# Decoding the covert shift of spatial attention from electroencephalographic signals permits reliable control of a brain-computer interface

**DOI:** 10.1101/2020.07.29.226456

**Authors:** Christoph Reichert, Stefan Dürschmid, Mandy V. Bartsch, Jens-Max Hopf, Hans-Jochen Heinze, Hermann Hinrichs

## Abstract

**Objective:** One of the main goals of brain-computer interfaces (BCI) is to restore communication abilities in patients. BCIs often use event-related potentials (ERPs) like the P300 which signals the presence of a target in a stream of stimuli. The P300 and related approaches, however, are inherently limited, as they require many stimulus presentations to obtain a usable control signal. Many approaches depend on gaze-direction to focus the target, which is also not a viable approach in many cases, because eye movements might be impaired in potential users. Here we report on a BCI that avoids both shortcomings by decoding spatial target information, independent of gaze shifts.

**Approach:** We present a new method to decode from the electroencephalogram (EEG) covert shifts of attention to one out of four targets simultaneously presented in the left and right visual field. The task is designed to evoke the N2pc component – a hemisphere lateralized response, elicited over the occipital scalp contralateral to the attended target. The decoding approach involves decoding of the N2pc based on data-driven estimation of spatial filters and a correlation measure.

**Main results:** Despite variability of decoding performance across subjects, 22 out of 24 subjects performed well above chance level. Six subjects even exceeded 80% (cross-validated: 89%) correct predictions in a four-class discrimination task. Hence, the single-trial N2pc proves to be a component that allows for reliable BCI control. An offline analysis of the EEG data with respect to their dependence on stimulation time and number of classes demonstrates that the present method is also a workable approach for two-class tasks.

**Significance:** Our method extends the range of strategies for gaze-independent BCI control. The proposed decoding approach has the potential to be efficient in similar applications intended to decode ERPs.

## 1. Introduction

Brain-Computer Interfaces (BCI) are designed to restore severely paralyzed people’s ability to communicate directly with their environment or to control assistive devices. However, most BCI systems are not suited for users with impaired eye movements since they require gaze shifts. Hence, several attempts have been made to implement BCI communication independent of eye movements. For instance, the modulation of oscillatory brain signals due to motor imagery is widely used to control a BCI [1, 2]. The shortcomings of motor imagery are a low number of distinguishable user intentions, high mental and training effort, and substantial performance variability among users. Accordingly, there is a high proportion of so-called BCI illiterates. A second group of gaze-independent BCIs are those that are controlled by stimulus-evoked brain activity modulations. One example is rapid serial visual presentation (RSVP), in which users watch a stream of varying symbols, sequentially presented at fixation. The detection of a target symbol is then decoded from an enhanced P300 response [3–5]. However, in RSVP tasks subjects are at risk to miss the target, due to fatigue during long stimulation sequences, as well as effects like attentional blinking [6]. A third approach is flicker stimulation, where visual stimuli flickering at selective frequencies are presented at different locations in the peripheral visual field. Shifts of attention to one or the other peripheral location are then decoded from power changes of the steady-state visual evoked potential (SSVEP) [7]. However, the accuracy of detecting SSVEPs from covert attention is largely reduced compared to overt attention. A fourth approach is to use auditory stimulation aiming at decoding event-related potentials (ERPs) [8, 9] or steady state responses [10]. But, auditory BCIs achieve considerably lower information transfer rates [11] and often still depend on vision to navigate through the system. An alternative channel to deliver stimuli is tactile stimulation, which has been used to decode BCI commands from P300 responses [12] and somatosensory steady-state evoked potentials [13]. In the latter study only 50% of participants could achieve control.

Most BCIs that use attention related brain activity as a control signal employ versions of the oddball paradigm and hence, focus on time as the feature that distinguishes the choice alternatives [14]. This typically requires long stimulation intervals, which renders the selection process time-inefficient and requires a high mental effort. An alternative brain mechanism suitable for BCI control is spatial attention, which is a particularly practicable approach when eye movements are impaired. That is, the spatial focus of attention can be covertly allocated to items in the visual field without the need to move the eyes [15, 16]. Importantly, a single stimulus can contain multiple target alternatives simultaneously, which allows for higher coding efficiency based on a comparably low number of trials. A prime stimulation paradigm would be the visual search task, where subjects focus attention to a target item among simultaneously presented nontarget items. Luck et al. [17] discovered that searching for a target elicits an electrophysiological response in visual cortex referred to as N2pc (N2 posterior contralateral). The N2pc is a lateralized negative polarity modulation of the ERP response in the N2 time range (180-300m) with stronger activity modulation over the posterior scalp contralateral to the target item. The N2pc therefore allows to assess a covertly attended item is located in the left or right visual field [18]. Numerous studies have investigated the N2pc, making it one of the best-characterized visual attention components in EEG and MEG (e.g., [17, 19–21]). Compared to the P300 that can be elicited by diverse target evaluation or categorization processes, the N2pc is a neural correlate very specific for visuo-spatial item selection. It shows a typical maximum at parieto-occipital electrode sites PO7/PO8. At a given electrode, the N2pc is derived by subtracting the response elicited by a target presented ipsilateral to the electrode from the response elicited by a target presented contralateral to the electrode [22]. There is firm evidence that the N2pc reflects the focusing of attention onto the target in visual search [19], and that it allows to track sequential shifts of attention [18]. The N2pc will be elicited as soon as subjects shift their attention to a target in the left or right hemifield. To maximize measured N2pc responses, the target should be easily detectable and presented among competing distractor items [21, 23–25]. Notably, spatial target selection processes can be derived for single stimulus arrays, i.e., there is no need for preceding presentations of non-target stimuli as typically used in oddball paradigms to elicit the P300. Most importantly, the N2pc is a comparably large amplitude modulation of the ERP, and therefore a very robust signal reliably appearing in single observers, which makes it an ideal tool to control BCIs.

Nonetheless, the number of studies investigating the N2pc as a control signal is low. Awni et al. [26] reported that averaging three trials for classification of the N2pc yields results comparable to other BCIs which implement covert attention. In a recent work it has been shown that an optimal subset of correlated components permits detection of target locations in single trials, superior to a common classification approach [27]. Single trial classification of visuospatial attention has also been investigated using hybrid N2pc and SSVEP features [28]. Another group tested single trial classification of the N2pc in aerial images [29], where they showed that the presence of an airplane and the visual hemifield in which it was present can be determined from the electroencephalogram (EEG). Fahrenfort et al. [30], found evidence that not only horizontally lateralized visual presentation but also presentation on the vertical midline can be decoded from EEG data using a linear discriminant classifier. While the aforementioned studies investigated the theoretical feasibility of the N2pc to control a BCI, there is no BCI implementation, in which users actively controlled the system and received feedback indicating the successful recognition of their attention shifts.

In the current study we present a closed-loop BCI that decodes which target color a user attended by determining the visual field (left/right) of its presentation from EEG signals. To discriminate four different colors, we presented all of them simultaneously in the visual search display but each color was associated with a unique sequence of left/right presentations, similarly to the principle in P300-based BCIs where each symbol is associated with a unique sequence of intensifications. The essential difference to P300 paradigms is that they depend on an oddball event, i.e. the time point of a rarely occurring event is determined from brain signals. Here, the target is present in every stimulus event, only differing in the visual hemifield where it is shown. Thus, some P300 BCIs might depend on the human’s ability to shift attention to peripheral locations but they decode temporal (present vs. absent target) rather than spatial differences in the brain response. Consequently, in matrix spellers the accuracy considerably decreases when subjects are not allowed to focus the target but the center of the matrix [31]. In contrast, our approach decodes the spatial differences (left vs. right) in an attention task because there are no stimuli where the target is absent. This makes the approach particularly useful for gaze-independent BCIs. We use a spatial filter approach that proved to be useful for classification of event-related potentials [32–34] and which we modified such that it allows us to discriminate the ERP modulations underlying the N2pc. We show that the features, extracted from training data and driving the BCI control, clearly represent characteristics of the N2pc, confirming that the BCI was driven by this marker of spatial attention.

## 2. Methods

### 2.1 Participants and Recordings

In this study 24 volunteers (15 female, mean age 27.8 years, σ=5.4 years) with normal or corrected to normal vision participated after they gave their written informed consent. The EEG was recorded in an acoustically shielded cabin, with the data being instantaneously processed to generate feedback from the decoded attention shifts. We recorded electroencephalographic activity at 29 electrode sites according to an extended ten-twenty system [35], referenced against the right mastoid using a BrainAmp DC amplifier and Ag/AgCl electrodes. The sampling rate was set to 250 Hz. For the online decoding we used only parietal and occipital electrodes (see figure 1), as the N2pc is known to show its topographical maximum at these sites [36]. Additionally, we recorded the horizontal electrooculogram (EOG) from electrodes at the outer canthi of both eyes and the vertical EOG from Fp2 and an electrode below the right eye for offline evaluation of eye movements. The study was approved by the local ethics committee of the Otto-von-Guericke University, Magdeburg and conducted in accordance with the principles embodied in the Declaration of Helsinki.

**Figure 1::**
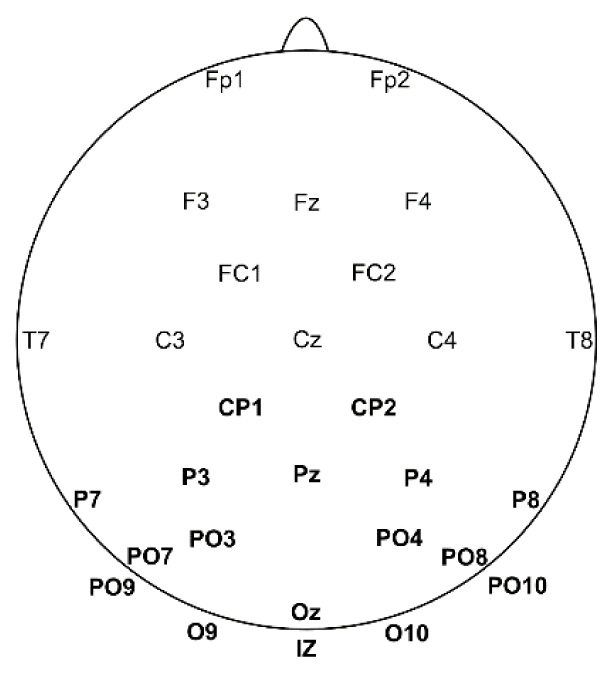
Layout of recorded EEG electrode positions (bold font indicates online processed channels).

### 2.2 Paradigm

The participants’ task was to navigate an abstract avatar towards a destination mark (a white cross). Figure 2 shows the temporal structure of a trial, required to achieve a brain-controlled movement step of the avatar. The movement direction of the avatar was controlled by shifting attention to a specific target item which randomly appeared in the left or right visual field. Each movement direction was associated with a specific color (blue/left, red/up, yellow/right, green/down), which was permanently visible at the margin of the playing field. The participants were instructed to fixate the avatar during the whole experiment and to prevent any eye movements except tracking the avatar after the end of the stimulation phase. At the beginning of every trial, the avatar changed its color for 2000ms to indicate the requested movement direction (e.g., red for moving the avatar up). Thereafter, a sequence of 12 visual stimuli started, in which item of all colors were presented simultaneously, either in the left or in the right visual field. The sequence was presented in pseudorandom order, such that the number of occurrences of pairwise colors on the same side was minimized and each color was presented six times in the left and six times in the right visual field. To focus attention onto the target colored item, participants were instructed to discriminate whether the ring with the cued target color had a gap, and they were told that a correct judgement would improve steering the avatar. The four items (colored rings), which formed a stimulus, were positioned at a horizontal visual angle of 4.26° and vertical visual angles of 1.64° and 2.73° relative to the avatar. Stimuli were shown for 250ms with an interstimulus interval of 500ms-750ms. As soon as all required EEG data were transferred to the BCI computer, the feedback was generated according to the decoding procedure described in section 2.4 and displayed for 2000ms, thereby giving the user feedback and time to prepare for the next trial. The feedback was presented by moving the avatar in the direction associated with the decoded color, plus showing emotional feedback (happy, neutral or sad facial expression) and by providing a sum of scores associated with the emotion (score +1: smiling face if decoding agreed with cued direction; score 0: neutral face if decoding did not agree with cued direction but the distance to the target cross was not increased by the movement; score −1: sad face if distance to the target cross was increased by the movement or the movement would imply a pass over the margin). If the avatar reached the destination mark, the face flickered for 2 s at 3 Hz, additional scores (number of the steps covered) were added and a new destination mark was positioned. We define a trial as the process of presenting the cue, directing attention to the target color while the stimulus sequence was presented and presenting the feedback (see Figure 2). After each trial we trained the classifier with the recently acquired data, i.e. we applied the decoder training as described in section 2.4. Several trials formed a run, in which subjects tried to achieve a maximal score. The run finished with the trial that exceeded a total run length of five minutes. Each participant performed eight runs, which amounted to an average total of 174 trials (σ =10).

**Figure 2:**
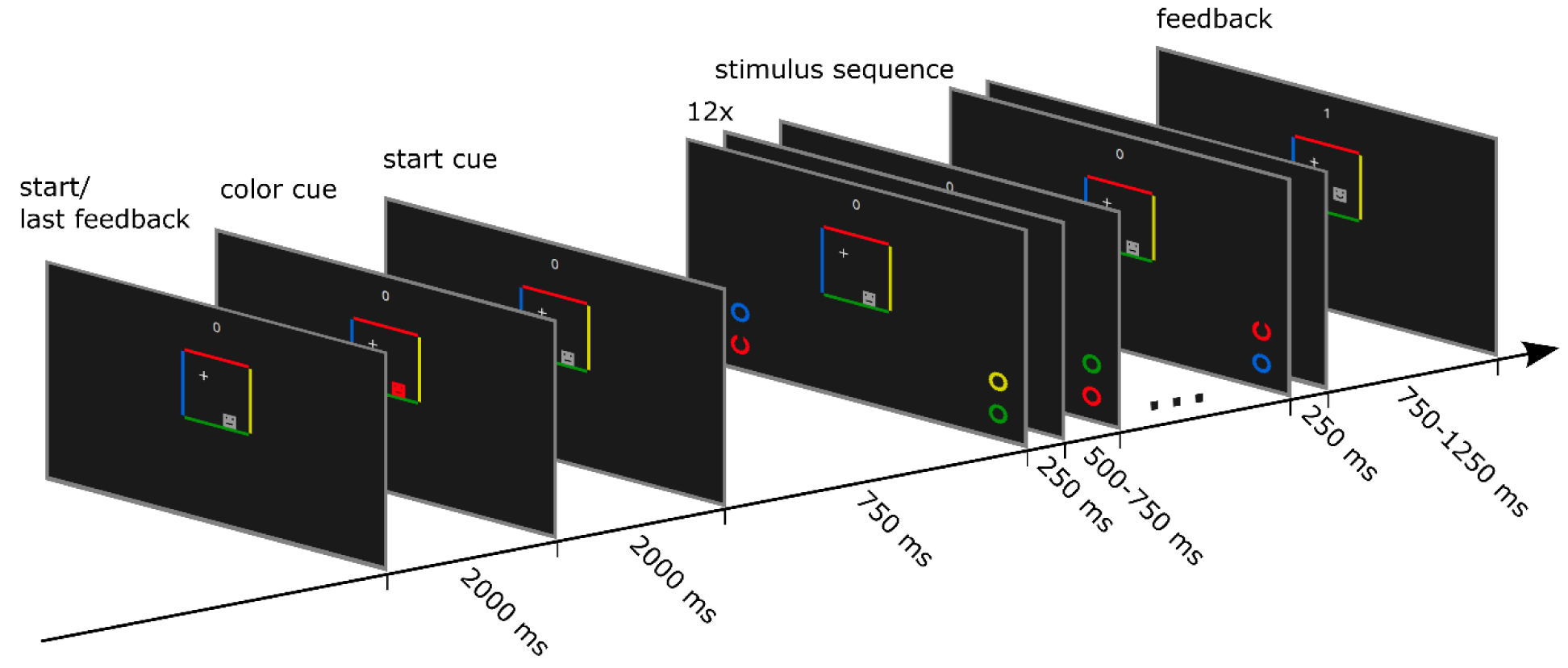
Structure of one trial. The direction of expected movement was cued by the respective color (here red for an upward move). During presentation of the stimulus sequence, subjects fixated the avatar but covertly directed their attention to the item (ring) of the cued color. Each colored ring switched the position in a pseudo-random but unique order such that the attended color can be determined from ERPs that are slightly differently evoked in the hemispheres and known as N2pc. The decoded color was fed back by a move of the avatar in the direction that accords to that color. A move in the cued direction was indicated by a smiling face and an increase of the score, an increase of the distance to the target cross was indicated by a sad face and a decrease of the score, and a move not cued but decreasing the distance to the target cross was indicated by a neutral face.

### 2.3 Stimulus Sequence

Each trial consisted of 12 stimulus onsets during which participants had to pay attention to the target drawn in the cued color. The decoding task was to detect whether an item in the left or in the right visual field was attended. The shape of the presented items could be either a ring or an upright Landolt-C. The presence of the gap was irrelevant for decoding the side of target presentation. It appeared randomly in a given trial and served to better focus spatial attention onto the precise location of the target. Since we aimed to discriminate four classes with only two conditions, on either side two potential targets were presented and hence, multiple stimulus combinations were required to determine the item the participant attended to. The stimulus sequence was pseudo-random such that i) each color appeared exactly six times at either side, ii) each possible pairing of the four colors appeared exactly eight times on opposite sides and four times on the same side. The top and bottom positions were also varied for each color but they were not relevant for decoding (see Table 1 for stimulus combinations). The rationale behind this approach is that each color is associated with a unique sequence of left/right presentations. We hypothesize that the lateralized differences in hemispheric ERPs correlate with the left/right presentations of the attended color. Thus, the aim of the decoder was to find the sequence of Left/Right presentations that fits the brain response best.

**Table 1:**
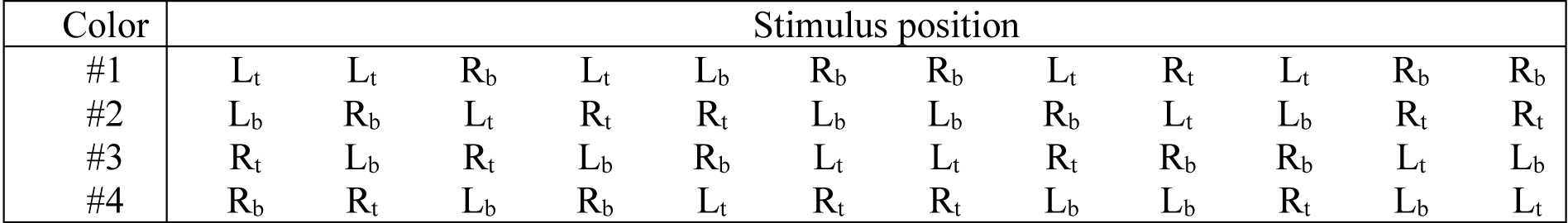
Sequence of stimulus item positions. Colors red, green, blue and yellow were randomly assigned to the position sequence of lines #1 to #4. Each column shows the positions (L_t_ – top left, L_b_ – bottom left, R_t_ – top right, R_b_ – bottom right) at one stimulus onset.

| Color | Stimulus position |  |  |  |  |  |  |  |  |  |  |  |
| --- | --- | --- | --- | --- | --- | --- | --- | --- | --- | --- | --- | --- |
| #1 | L<sub>t</sub> | L<sub>t</sub> | R<sub>b</sub> | L<sub>t</sub> | L<sub>b</sub> | R<sub>b</sub> | R<sub>b</sub> | L<sub>t</sub> | R<sub>t</sub> | L<sub>t</sub> | R<sub>b</sub> | R<sub>b</sub> |
| #2 | L<sub>b</sub> | R<sub>b</sub> | L<sub>t</sub> | R<sub>t</sub> | R<sub>t</sub> | L<sub>b</sub> | L<sub>b</sub> | R<sub>b</sub> | L<sub>t</sub> | L<sub>b</sub> | R<sub>t</sub> | R<sub>t</sub> |
| #3 | R<sub>t</sub> | L<sub>b</sub> | R<sub>t</sub> | L<sub>b</sub> | R<sub>b</sub> | L<sub>t</sub> | L<sub>t</sub> | R<sub>t</sub> | R<sub>b</sub> | R<sub>b</sub> | L<sub>t</sub> | L<sub>b</sub> |
| #4 | R<sub>b</sub> | R<sub>t</sub> | L<sub>b</sub> | R<sub>b</sub> | L<sub>t</sub> | R<sub>t</sub> | R<sub>t</sub> | L<sub>b</sub> | L<sub>b</sub> | R<sub>t</sub> | L<sub>b</sub> | L<sub>t</sub> |

### 2.4 Decoding Algorithm

All data processing was performed using MATLAB® R2016a. In order to determine the most probably attended sequence, we used a statistical learning technique that autonomously estimated the optimal spatial weighting of channels and simultaneously estimated the time course of event-related potentials in surrogate channels. We first introduced a variant of this method in [37] and verified its efficiency in a P300-based closed-loop BCI in [33]. After a stimulation phase was finished, we treated the interval starting from the first stimulus and ending at the start of the feedback phase as one trial by cutting out that segment of raw EEG data. Subsequently, we band-pass filtered the data from 1.0Hz to 12.5Hz using an 8th order Butterworth zero-phase digital IIR filter (*filtfilt* function in MATLAB®) and resampled the data with a 50Hz sampling rate to reduce the amount of data and to remove redundancy.

Let these preprocessed data be a matrix ***X*** ∈ ℝ^*n×m*^ with *m* the number of channels and *n* the number of samples. For each potential target color *c* we modeled a reference function ***Y***^*c*^ ∈ ℝ^*n*×*k*^, where *k* is the number of samples within the processing interval following a single stimulus onset. Here we chose a processing interval length of 720ms which resulted in *k* = 36 samples. Each column *i* in *Y*^*c*^ represents a reference function, modeling the *i*^*th*^ sampling point after stimulus onset *t*_*s*_. We set 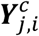 to if *j* = *t*_*s*_ + *i* and the potential target *c* appeared in the left visual field, to −1 if *j* = *t*_*s*_ + *i* and the potential target *c* appeared in the right visual field and to 0 otherwise. In other words, we have embedded into ***Y***^*c*^ the identity matrix *I*_*k*_ following a presentation of *c* on the left and -***I***_*k*_ following a presentation on the right. With the matrix multiplication

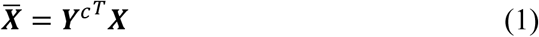

***Y***^*c*^ can be considered to be a weighting matrix that subtracts the sum across stimuli in ***X*** representing right visual field stimulation from the sum across stimuli in ***X*** representing left visual field stimulation and thus, 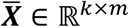 represents the difference of pooled brain data following left presentations and pooled brain data following right presentations. This is essentially what is typically done when isolating the N2pc component, i.e. considering difference waves [18, 20]. While determining the mean difference per channel is sufficient for grand mean analysis, different steps are required to classify ERPs in short EEG recordings. Here we use canonical correlation analysis (CCA) to estimate a set of coefficients serving as spatial filter from a set of training data:

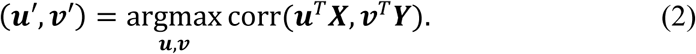

Here, EEG data of all trials are concatenated in ***X*** and the corresponding reference functions of the known target color are concatenated in ***Y***. The coefficients in ***u***^′^ can be considered to represent a spatial filter that combine the channels to a surrogate channel while coefficients in ***v***^′^ modulate the waveform that maximally correlates with that surrogate channel. CCA reveals min{*m, k*} components where the canonical correlation coefficient is decreasing with each component. This approach reduces dimensionality and noise, is data-driven and transforms EEG signals such that we can determine the attended sequence by an ordinary correlation measure. Note that the weightings in ***Y*** enable the CCA to perform optimization subject to the difference waves rather than ERPs. Thus, small modulations of ERPs can be isolated as it is done in typical N2pc studies.

In order to estimate ***u***^′^ and^′^, we used a set of training data where *Y* was modeled according to the sequence of stimulus positions assigned to the target color the participant attended during acquisition of data in ***X***. We used the MATLAB® function *canoncorr* of the Statistics and Machine Learning Toolbox™ to solve Equation 2. This procedure we refer to as decoder training. It is applied online after each trial using all training data available and offline in every step of cross-validation using the training data set.

Once the canonical coefficients are determined, they can be used as spatial filter on new data. Given new data in matrix ***X*** obtained from a new trial, we calculated the correlation *ρ* = corr(***u***′^*T*^***X, v***′^*T*^*Y*^*c*^) for each color *C*. The color that yielded the highest average correlation with the first two components was decoded as the attended color. This procedure we refer to as decoding. It is applied online after each presentation of a stimulation sequence to determine the feedback to be presented and in offline analyses in every step of cross-validation using the left-out test data set. See figure 3 for an example, how the target color is determined from stimulus sequences.

**Figure 3:**
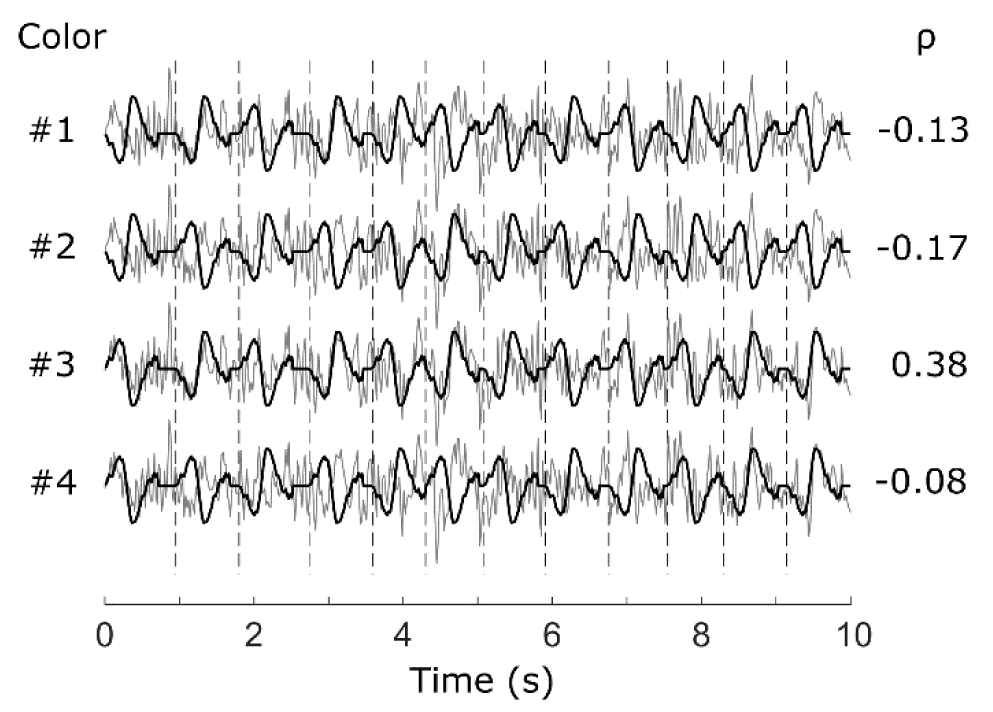
Decoding example. The gray lines represent the surrogate channel signal ***u***′^*T*^***X*** during stimulation which is compared with each color-related signal ***v***′^*T*^*Y*^*c*^ (black lines). Vertical dashed lines denote the onset of a stimulus. Each color is associated to a sequence of left/right presentations, which are modeled in *Y*^*c*^ and are reflected by representations of ***v***′ for left and -***v***^′^ for right presentations in ***v***′^*T*^*Y*^*c*^. For each color the correlation ρ = corr(***u***′^*T*^***X, v***′^*T*^*Y*^*c*^) is calculated. The highest correlation coefficient indicates the sequence, most similar to the brain signals (here #3) and is selected as target color.

### 2.5 Evaluation

In offline analyses we used leave-one-run-out cross-validation where decoder training was performed with data from seven runs and decoding was performed on the left-out run. We calculated the decoding accuracy as the percentage of correctly decoded trials. The chance level and confidence interval were determined by means of permutation testing. For this purpose, we permuted the labels that assign the target color to the EEG data across trials and performed a cross validation analysis with this data set. To estimate a confidence interval for guessing, we repeated this permutation 1000 times for each participant and fitted the achieved decoding accuracies to a normal cumulative distribution function.

For statistical comparisons between two sets of individual decoding accuracies we performed a Wilcoxon signed-rank test, unless stated otherwise.

As a measure of the trade-off between accuracy and speed of a system, the information transfer rate (ITR) has been established in BCI research. We calculated this measure according to [38] and report it where appropriate.

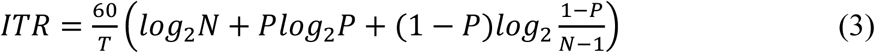

where *N* is the number of trials, *P* is the decoding accuracy and *T* is the trial duration in seconds, yields the ITR in bit/min.

### 2.6 Contribution of ERPs

In typical ERP studies including most N2pc studies, the grand average ERP is calculated from a group of subjects without considering individual differences and involving all trials which showed behavioral evidence that the search task was correctly processed. Here we compare this common approach with a single trial approach where we distinguish between trials that could be correctly predicted in a single-trial classification and trials that could not correctly be predicted, to uncover the cause of potentially failing BCI control. Specifically, we applied the decoding algorithm in a cross-validation scheme to single stimulus presentations, in order to discriminate between targets presented in the left versus the right visual field. Note that the determination of canonical components is identical with that in the sequence decoding approach because the modeling of the reference functions reveals the same matrices ***X*** and *Y*. In the left-out data, the correlations ρ = corr(***u***′^*T*^***X, v***′^*T*^*Y*) of the first two canonical components were averaged. This measure gauges the similarity of brain signals to left and right target presentations, respectively. Trials yielding a negative correlation were classified as left target presentation, and trials with a positive correlation as right target presentation. Depending on the individual decoding accuracy, we determined subsets of trials representing confident and unconfident predictions. Specifically, we kept half of all correctly predicted trials as confident predictions and rejected those where |ρ| was low. Likewise, we kept half of all incorrectly predicted trials as unconfident predictions and rejected those where |ρ| was low. This approach discards trials with highest uncertainty of the learning algorithm’s decision. Using the subsets of confident predictions and unconfident predictions, we calculated grand average data for the whole group of subjects to reveal the difference between correctly and incorrectly decoded brain signals. Furthermore, we determined the canonical components of the whole group to find out which features are generally used by our data-driven decoding approach.

## 3 Results

### 3.1 Decoding performance

Most participants (22 out of 24) generated online control commands with an accuracy higher than 55% (chance level: 25%). The average decoding accuracy was 68.4% (σ=16.7%). Half of the participants achieved more than 70% correctly decoded commands (figure 4(a)). The critical accuracy level of 70% is considered to be required to effectively control a BCI [39, 40]. Note that for online decoding the training data set was increased with every new trial. In contrast, in offline decoding we can use much more training data when performing cross-validation, revealing a better estimate of the long-term performance. Therefore, we performed leave-one-run-out cross-validations in all subsequently reported analyses.

**Figure 4:**
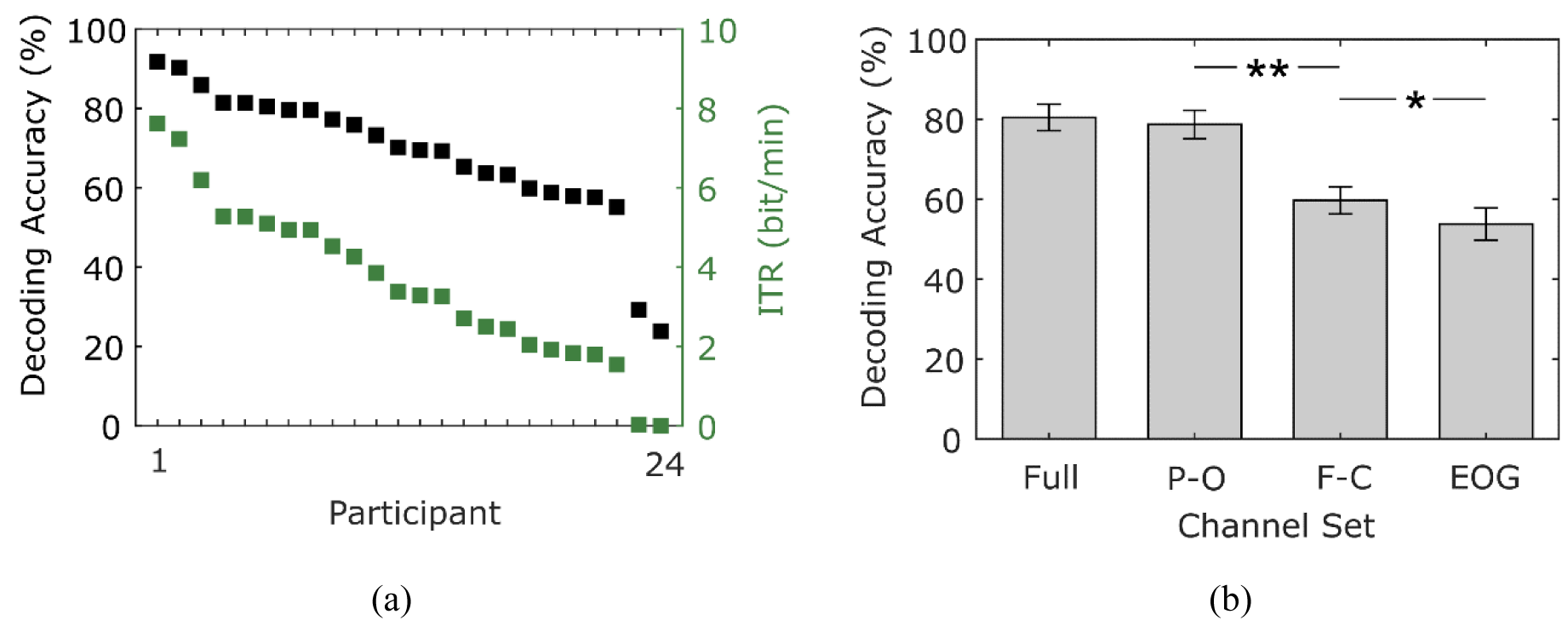
(a) Decoding accuracy achieved online (black) for single participants (sorted by online accuracy) and the corresponding ITR (green). (b) Average decoding accuracy and standard error of the mean achieved in offline analysis using different channel sets.

We performed permutation tests using the same channels as we used for the feedback generation during the experiment. The permutation tests revealed a mean chance level of 25.0% (σ=0.17%), an upper 95% confidence interval of 31.9% (σ=0.39%) and an upper 99.9% confidence interval of 38.0% (σ=0.68%). This implies that decoding accuracies higher than 38% can be considered significantly different (p<0.001) from a guessing classifier.

Using our data-driven decoding algorithm, we compared different sets of channels to evaluate the performance of the parieto-occipital (PO) channel set (used online, bold labels in figure 1), the fronto-central (FC) channels (not used online), the full channel set, as well as EOG only. With the cross-validation approach the average decoding accuracy increases significantly (p<0.001) from 68.4% to 78.7% (σ=17.4%) using the PO channel set, compared to the online performance. The higher number of training samples, which permits a better estimate of a well-trained BCI, results in a larger number of subjects (N=22) who could efficiently control a BCI (>70% accuracy). Importantly, the FC channels which we excluded from the online experiment showed a significantly reduced performance compared to the PO channels (59.7% compared to 78.7%, p<0.001), although decoding accuracy was significantly above chance level for 21 participants. This implies, that the fronto-central electrodes capture informative signals to decode the lateralized occurrence of the target item. The decoding accuracies achieved with only EOG channels further decreased significantly (p<0.05), but the relatively high performance of 53.8% (σ=19.8%; 17 participants above chance level) suggests that eye movements might play a role to some extent in controlling the BCI. However, the decoding accuracies of single subjects did not correlate significantly between EOG and PO channel sets (ρ=0.21) indicating that the decoder did not benefit from information related to eye movements when using only PO channels.

All previous analyses were performed using the full sequence of 12 stimuli presented in one trial. To investigate the accuracy achievable with less stimuli and consequentially in shorter time, we tested our decoding algorithm with sequence lengths of 1 to 12 stimuli per trial and calculated the corresponding ITR. While the decoding accuracy consistently increased (p<0.05, Bonferroni-corrected) with every additional stimulus, the ITR was already maximal with 6 stimuli (see figure 5(a)), suggesting that the highest information flow could be achieved in half of the time (average ITR: 5.9 bit/min, σ=3.1 bit/min) but with an average accuracy of only 64.7% (σ=31.8%). We performed the same analysis with a two-class decoder in order to investigate the performance of our approach for binary control. Note that the chance level in this case is at 50% and not 25% as with four classes. In this analysis we assumed that there are only two colors, one target color and one nontarget color. This is possible with our data since only the side of target presentation matters and not the number of distractors presented. The results of this analysis are shown in figure 5(b). Again, the decoding accuracy is consistently increasing with longer stimulation intervals, achieving an average of 90.7% (σ=12.7%) at the maximum number of 12 stimuli. Here, the highest average ITR of 5.3 bit/min (σ=2.5 bit/min) is reached with 4 stimuli where the average accuracy is 80.6% (σ=11.4%). Notably, 19 out of 24 participants achieve decoding accuracies exceeding 90% with the full stimulation sequence and 10 participants even with half of the stimulation sequence.

**Figure 5:**
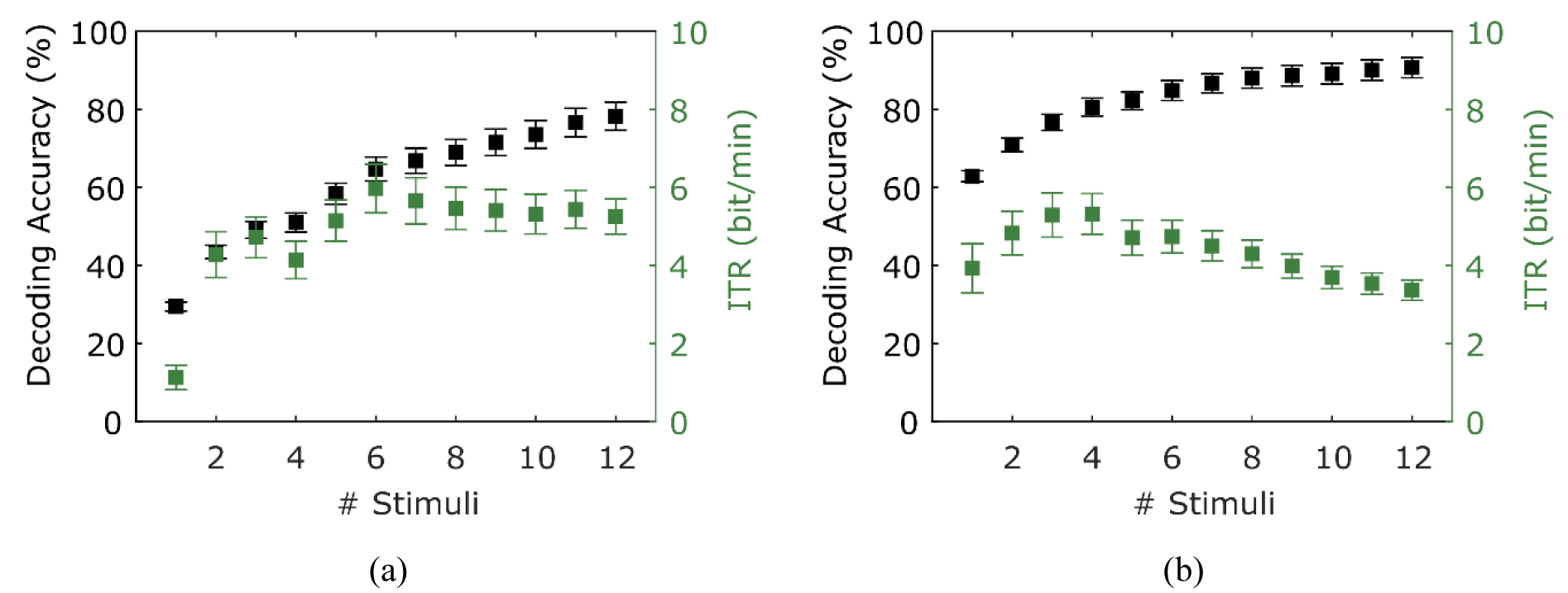
Average decoding accuracy (black) and corresponding ITR (green) depending on the number of stimuli involved in the decoding. (a) shows the results of a four-class decoder (chance level = 25%; note that a nonambiguous recognition is not possible with only one stimulus) and (b) shows the results of a two-class decoder (chance level = 50%). Error bars indicate standard error of the mean.

### 3.2 Waveform analysis

Next, we examined which features of the ERP response contributed most to the decoding of the target lateralization, and hence affected the control of the BCI. Our decoding approach is based on the correlation values of the first two canonical components, determined from a set of training data. Due to the positive and negative weightings in the reference functions, related to left and right target presentation, the coefficients in ***v***′ represent a kind of canonical difference wave. Thus, the waveform of the canonical component ***u***′ defines the time course of the signal at a surrogate channel as determined with CCA, maximally correlating with the canonical difference wave ***v***′.

Here we consider the canonical components and difference waves of left and right visual field presentation for the whole group of subjects. We compare the involvement of all trials with involvement of only confident predictions, i.e., the trials we considered, according to our description in (2.6), as correctly predicted (22% of all trials) and falsely predicted (27.5% of all trials), respectively (figure 6). Figure 6(b) shows the first two canonical components, scaled by their canonical correlation. Because the correlation is scale invariant, the units of canonical components are arbitrary. The first component correlates highly with the difference wave in channel PO7 showing a positive peak around 250ms. The second component correlates with the difference wave in channel PO8 showing a considerably weaker, negative peak around 230ms. The N2pc is represented as positive deflection at PO7 because we show the difference of right presentations subtracted from left presentations and thus, contralateral activity is subtracted from ipsilateral activity. In contrast, at PO8 a negative deflection appears because of subtracting ipsilateral from contralateral activity. The localization, timing and stronger expression over the left hemisphere are characteristic for the N2pc component. Note that we have used the PO channel set to determine the canonical components and decode the side of target presentation, but show averages of all channels in the topographic maps.

**Figure 6:**
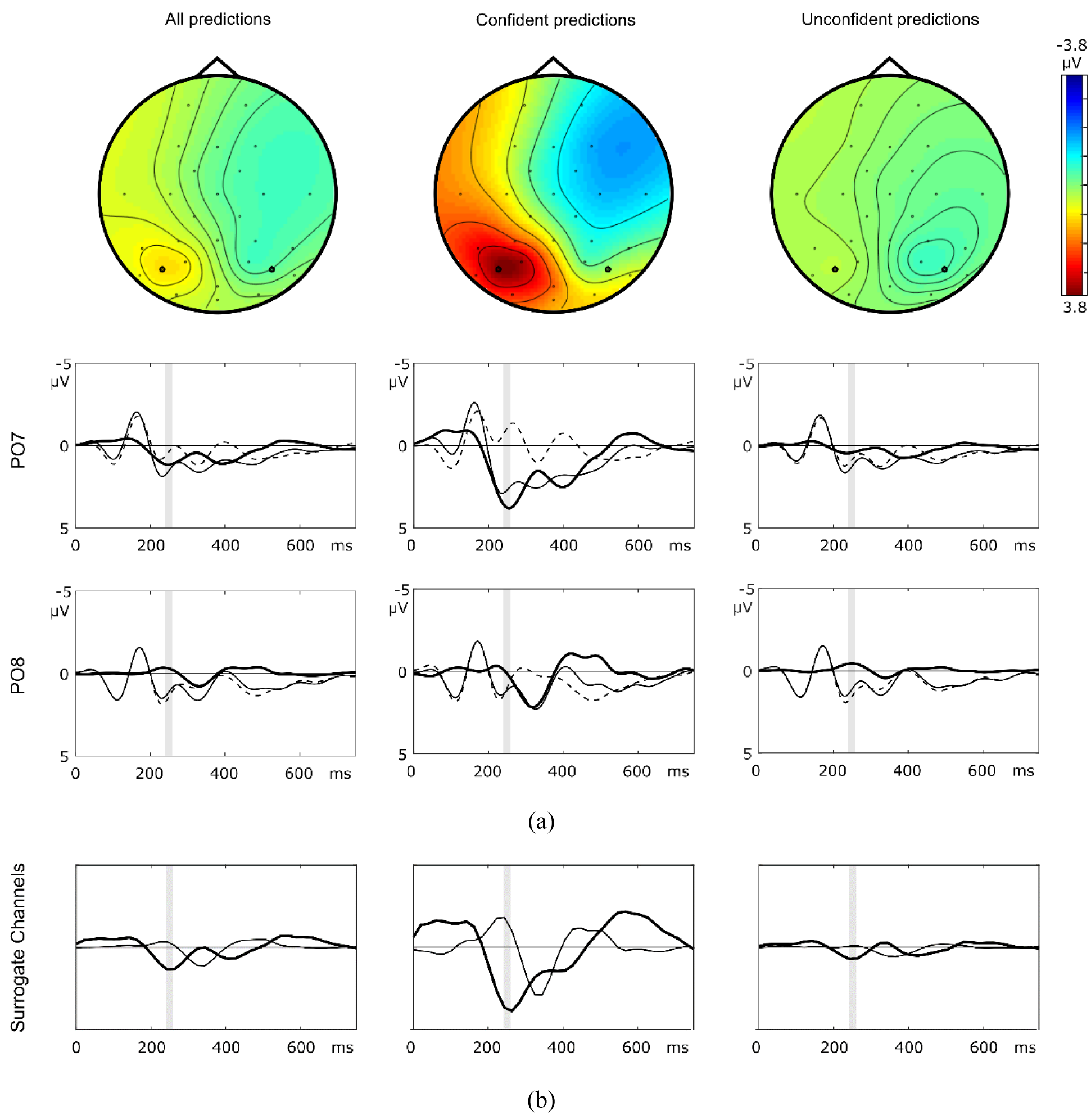
(a) Topographic maps of the grand mean difference wave (right target presentation subtracted from left target presentation) in the interval 240-260ms after stimulus onset calculated across all trials (left column), only trials yielding confident predictions (central column, 22% of all trials) and only trials yielding less confident predictions (right column, 27.5% of all trials). Diagrams below the topographic maps show the time course at channels PO7 and PO8 for target left (solid lines) target right (dashed lines) and the difference of both (bold lines). The shaded area denotes the interval used to calculate topographic maps. (b) Surrogate channels as determined by CCA. The first canonical component (bold lines) highly correlates with the difference wave at PO7, the second canonical component (solid line) highly correlates with that of PO8 except in the case of unconfident predictions.

As an important result, we find that when considering only trials which can be confidently classified as left or right target presentation, the N2pc amplitude is considerably increased. In contrast, when taking into account only incorrectly classified trials, the N2pc is reduced at PO7 compared to including all trials. Also, while successfully predicted trials show a positive peak at PO7 around 400ms and at PO8 around 320ms, these effects are much weaker in wrongly predicted trials. This suggests that successful decoding of attention directed to a target presented in the left or right visual field is mainly accounted for by an ERP modulation reflecting the N2pc.

### 3.3 Interindividual variability

The typical group-averaged N2pc is a lateralized negativity over the hemisphere contralateral to the target visual field. It often displays a stronger lateralization to one hemisphere (usually to the left). The lateralization varies considerably among subjects and sometimes even a right-sided lateralization is observed [20]. Here, we differentiate between left and right lateralized brain patterns, based on a subset of correctly predicted trials. This demonstrates the individual variability of brain signals, which is important for subject-specific BCI control. To demonstrate the lateralization variability, we selected subjects showing a higher absolute difference wave at PO7 in the interval 240-260ms compared to PO8 (n=19) and vice versa (n=5). The resulting group-specific grand averages are shown in figure 7. Due to the much higher number of subjects in the left lateralized group the pattern is quite similar to that of the grand average. The right lateralized group shows a difference wave at PO8 negatively correlating with that of PO7 in the left lateralized group. While such individual differences of overall lateralization cancel out in typical grand average ERP analyses, they are important for BCI control which must rely on individually trained learning algorithms.

**Figure 7:**
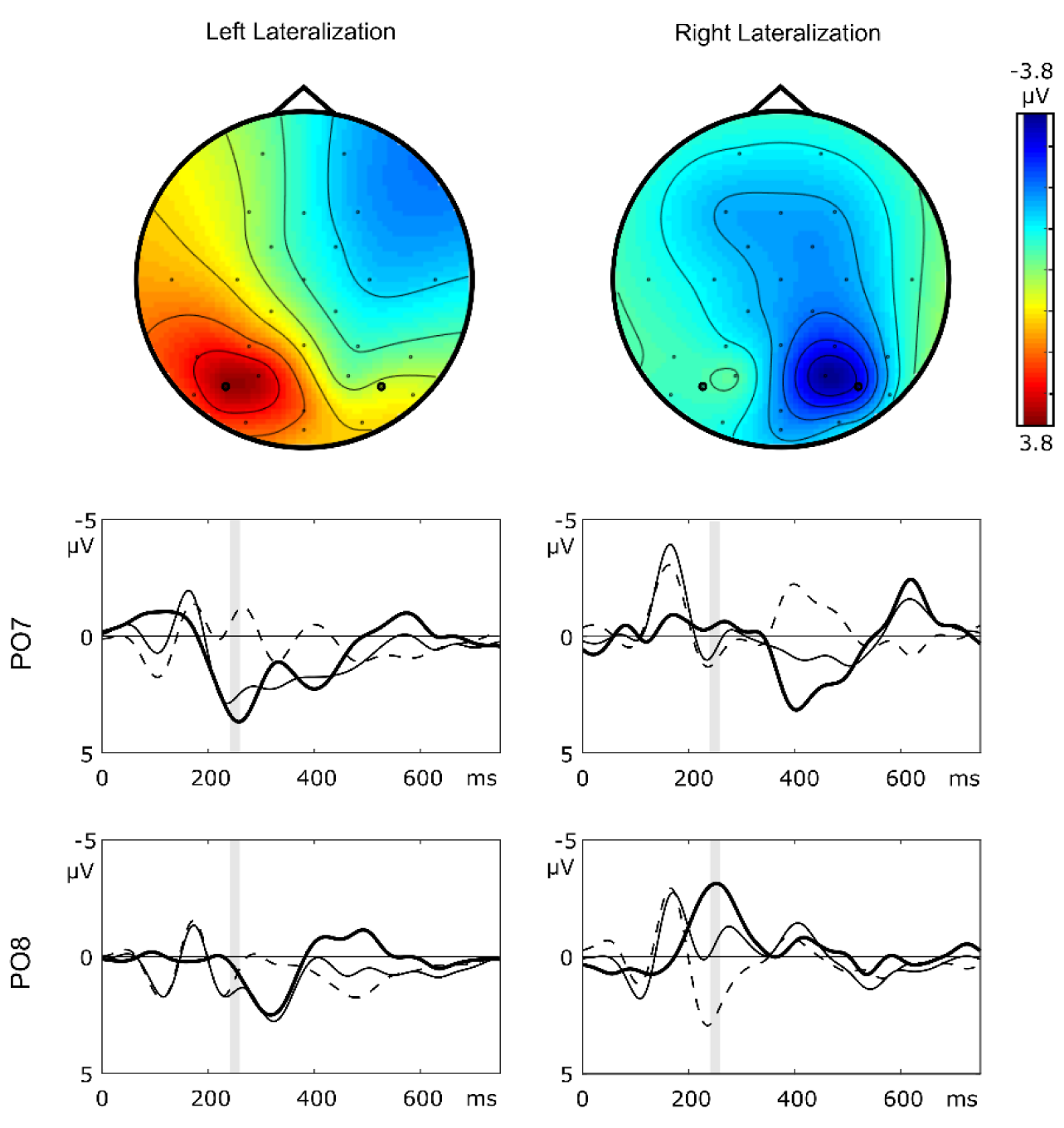
Topographic maps of the grand mean difference wave in the interval 240-260ms after stimulus onset calculated across subjects showing the N2pc preferably at the left hemisphere (left column) versus the right hemisphere (right column). The time-amplitude diagrams below the topographic maps show the ERP waveforms at channels PO7 and PO8 for left (solid lines) and right targets (dashed lines) together with the corresponding difference-waves (bold lines). The shaded area denotes the interval used to calculate topographic maps.

## 4. Discussion

The N2pc is a reliable marker of shifting attention to a target in visual search, but whether the N2pc is robustly elicited at a single trial level is not investigated so far. In this study, we show that the shift of spatial attention as indexed by the N2pc can reliably be decoded from a short series of single stimulus-evoked brain responses. Its robustness even allows to control a BCI. So far, the N2pc component has not been established in BCI applications, despite the fact that it is well suited as a control signal, because it does not rely on eye movements. Most attention-based BCIs used the P300 response, elicited by oddball stimuli [14]. The main difference to commonly used paradigms using visual attention is that in the present approach, target and non-target stimuli are presented at the same time on every stimulus onset, differing only in terms of spatial location. The present approach merely relies on covert shifts of spatial attention. It is, therefore, independent of gaze shifts – a feature that is central for BCIs aiding patients with gaze paralyses.

The traditional N2pc is a grand average effect, derived by contrasting trials with attention shifts to left versus right visual field targets. So far, it is not sufficiently clear, whether the N2pc would be a reliable enough signal that appears at a single trial level. Only a few studies aimed at detecting the N2pc in single trials. In the BCI study proposed in [29] no distractor was presented in the opposite visual field and thus the lateralized brain activity may simply reflect low-level physical stimulus lateralization which may capture attention in a bottom-up fashion. This is in contrast to balanced search arrays, typically used to elicit the N2pc, where top-down signals solely control the shift of attention to the target [19]. Thus, the approach in [29] would not be suited to discriminate between choice options as it confounds decoding with signals reflecting physical lateralization. In contrast, the study of Fahrenfort et al. [30], used a top-down approach and achieved decoding accuracies slightly above 60% correct in a sample-by-sample two class decoding approach. This is comparable to the present results when using only one stimulus in the two-class decoder analysis with our data. However, involving several repetitions of single stimuli increased the accuracy of the decoder in line with findings of Awni et al. [26] who averaged up to 3 stimuli. We demonstrated that a single stimulus elicits a measurable contralateral negativity that would combine over trials to result in the known grand average N2pc. We showed that the components extracted by the spatial filter approach and inducing the classification correlate with the characteristics of the N2pc, confirming our hypothesis that brain activity reflecting visual spatial attention is the main driver in this BCI. To further investigate this issue, we compared single trials with a high likelihood that predictions were confident with those with a low likelihood. This analysis clearly showed that the grand average N2pc effect is much higher in trials decoded correctly with high confidence but weak when predictions were wrong. This finding first corroborates our hypothesis that the BCI is mediated by brain signals reflecting spatial attention. Second, it suggests that subjects fail to control the BCI, when hemispheric differences are not specifically enough reflected in single trial EEG data.

The typical N2pc is derived by averaging across subjects, trials and even across both hemispheres [18, 41, 42], which neglects individual differences in activity lateralization. To control BCIs efficiently, individually trained classifiers are essential, because transfer learning across subjects commonly results in a substantially reduced accuracy of the decoder [33, 43, 44]. When calculating the grand average in our study, the voltage distribution of the N2pc peak showed a lateralization to the left hemisphere, due to the majority of subjects (n=19) showing such left lateralization. Previous grand-average analyses of the N2pc in the MEG observed a lateralization to the left hemisphere as well, but also showed individual hemispheric dominance at a source level [20]. The N2pc has also been found to be confined to the left hemisphere when targets and nontargets were words of different semantic categories [19]. Lateralization is often observed in attention-related brain processes. For example, when targets are laterally presented in a dual stream using RSVP, a left visual field advantage has been reported, as indicated by better identification of targets, an earlier N2pc, and higher P300 amplitudes in the right hemisphere [45].

While our approach was initially designed to decode four different commands, the attention shift to the left and right visual field is most suitable for binary decoding. In fact, a simulation of binary decoding using our recorded data revealed high accuracies exceeding those of other binary BCIs using control strategies independent of eye movements such as motor imagery [46] or attention shifts to auditory stimuli [9].

We implemented a closed-loop BCI, which by design can only be trained on past trials, and provided feedback from the first trial. In this mode, half of the subjects did not exceed the critical level of 70%, which is assumed to allow for effective BCI control [39, 40] although most of them performed clearly above chance. However, when more training data were provided as it is the case in cross-validation and in long-term BCI use, all except two BCI illiterates exceeded the effective BCI control level. In contrast to previous studies our BCI-illiteracy rate is comparably low. Only 8% of participants were not able to control the BCI. BCI-illiteracy is an often-reported issue frequently observed in BCIs that are based on motor imagery [47] and amount to a total of 15%-30% of the population [48]. In this respect our study excels other approaches. The reason why BCI control is not achievable in some people is still debated. Differences in anatomical structures of individual brains are most likely to play a major role [49] giving rise to a high variability of imagery in the population [50]. While individual differences are often not in the focus and averaged out in studies targeting group level effects, they have to be taken into account in BCI setups. Here, classifiers are trained on individual brain signals, resulting in a distribution of performance over subjects.

To generate feedback, we decoded spatial attention from parieto-occipital channels, where the N2pc typically shows a maximum [51]. We used a spatial filtering algorithm which determines channel contributions from a given channel set and thereby considers individual differences in the evoked brain patterns, preventing the requirement of elaborate channel selection strategies [52, 53]. In offline analyses we also evaluated the performance of the fronto-central channels and the EOG channels, which were not used for online decoding. Although the number of errors was significantly higher compared to misclassifications revealed by the parieto-occipital channel set, the achieved accuracies were above chance, potentially due to eye-movements and activity in frontal cortex contributing to the side-decoding. Given that the present study was performed in healthy subjects without impairments of eye-movements, unintentional small saccades might have accompanied shifts of spatial attention. Nevertheless, the mere fact that the parieto-occipital channels provide the highest decoding accuracy, which did not correlate with the accuracy achieved with the EOG across subjects, indicates that the features that drive the discrimination of lateralized target presentation originate from the visual cortex areas consistent with source localization analyses of the N2pc [20].

An accepted metric to validate BCI performance is the information transfer rate, which represents a measure of the accuracy/speed tradeoff. We investigated how accuracy and ITR changes with the number of stimuli and consequently with the time required to induce a command. While accuracy improved with a higher number of stimuli, the ITR showed a maximum at half of the actual presentation time in four class decoding and at even shorter presentation time in two class decoding. Given these results we would opt for more reliable decoding at longer stimulation time rather than maximizing the ITR by accepting more errors.

We consider the ITR to be a suboptimal metric for validating BCIs because its baseline value corresponds to the chance level and not to the level of effective control. Furthermore, if short classification intervals are used and the time the brain needs to process the feedback is ignored, the ITR might be overestimated. For example, Xu et.al. [28] tested a hybrid SSVEP/N2pc paradigm and calculated an ITR of 23.6 bit/min, assuming that the binary classification with an accuracy of 72.9% can be performed every 400 ms. They compared their result with 15 other studies on gaze-independent BCIs of which 8 achieved ITRs exceeding our results. In the review by Riccio et al. [11], where several gaze-independent BCIs were listed, 8 of the 37 reported ITRs were higher than in our study. Only one of these BCIs used visual spatial attention as discriminative feature, which was determined by SSVEPs and achieved only 0.91 bit/min on average [54]. Gaze-independent BCIs implementing an RSVP spelling system have been reported to approach ITRs up to 20.3 bit/min [3, 5]. However, in these studies up to 300 stimuli were presented at a fast pace which requires a high cognitive load to perceive the target. The visual search approach we presented here could therefore serve as an alternative to RSVP as it requires lower mental effort and might achieve improved performance when presentation parameters are optimized in future research.

### Conclusion

We presented a new method to control a BCI focusing exclusively on spatial attention, independent of gaze shifts using ERP decoding of the N2pc response. The results show that the N2pc can be reliably decoded from short stimulus sequences, which renders our method a time-efficient approach for attaining highly accurate discrimination of multiple commands and even higher accuracy for binary decisions. Future work will focus on a simplification of the stimulus design (larger items, less choice alternatives), to make it suitable for locked-in patients suffering from low vision. The here proposed N2pc controlled BCI could become a superior alternative to BCIs that fail in patients with impaired eye movements and a limited capability to perform long stimulation sessions.

## Acknowledgements

The authors declare that there is no conflict of interest regarding the publication of this article. This work was partially funded by the Federal Ministry of Education and Research, Germany, grant number 13GW0095D.

## Notes

### Competing Interest Statement

The authors have declared no competing interest.

